# Multicellular, IVT-derived, unmodified human transcriptome for nanopore-direct RNA analysis

**DOI:** 10.1101/2023.04.06.535889

**Authors:** Caroline A. McCormick, Stuart Akeson, Sepideh Tavakoli, Dylan Bloch, Isabel N. Klink, Miten Jain, Sara H. Rouhanifard

## Abstract

Nanopore direct RNA sequencing (DRS) enables measurements of RNA modifications. Modification-free transcripts are a practical and targeted control for DRS, providing a baseline measurement for canonical nucleotides within a matched and biologically derived sequence context. However, these controls can be challenging to generate and carry nanopore-specific nuances that can impact analysis. We produced DRS datasets using modification-free transcripts from *in vitro* transcription (IVT) of cDNA from six immortalized human cell lines. We characterized variation across cell lines and demonstrated how these may be interpreted. These data will serve as a versatile control and resource to the community for RNA modification analysis of human transcripts.

## Context

Nanopore Direct RNA sequencing (DRS) has emerged as a method for analyzing native RNA strands based on ionic current disruptions during translocation through a biological pore. Deviations in the ionic current disruptions may be attributed to RNA modifications[1], such as pseudouridine(Ѱ)[2]*, N*^6^-methyladenosine (m^6^A)[3], and inosine (I)[4]. Oxford Nanopore Technologies has developed basecallers capable of identifying modifications in RNA and DNA but are limited in scope. Out of the 170+ known RNA modifications[5], only m^6^A can be identified *de novo* with a basecaller[6]. For other modifications, negative controls and additional bioinformatic tooling are necessary. These negative controls include but are not limited to enzymatic knockouts of RNA modification machinery[7], synthetic RNA controls[8–10], expected distribution pore models for computational analysis[11,12], and genomic and transcriptomic templates for *in vitro* transcription (IVT)-based negative controls (i.e., unmodified transcriptomes)[2,13].

IVT-derived, unmodified transcriptomes[13,14] are an attractive option for analyzing modifications using DRS[2,13,15–17]. Currently, m^6^A is the only exception with several signal-focused tools, including Dorado[18], the Oxford Nanopore RNA base caller, and m6anet[6], a multiple instances learning-based neural network. To generate these IVT datasets, polyadenylated (poly-A) RNA strands are reverse transcribed to cDNA, PCR amplified, and then *in-vitro* transcribed into RNA using canonical nucleotides. This process maintains the sequence context of the initial poly(A) RNA sample while “erasing” the RNA modifications, providing a baseline to compare putative modification sites. However, IVT RNA derived from a single cell line may not comprehensively capture the landscape of expressed genes in the human transcriptome, for example, if applied to a different cell line. IVT RNA derived from multiple human cell lines could better capture these differences and be applied broadly.

We present a long-read, multicellular, poly-A RNA-based, IVT-derived, unmodified transcriptome dataset for DRS modification analysis. We identified and flagged positions where the IVT data set differs from the GRCh38 reference, which could result in the false identification of modification sites. We also propose a strategy for filtering out these sites. This includes several mismatch tolerance levels that an end user can select. We also created a pooled version of this IVT dataset for increased representation of genes and positions of interest at the cost of cell line specificity. Finally, we computed ionic current-level alignments for each cell line, allowing users to apply this dataset without additional preprocessing steps.

This publicly available dataset will be a resource to the direct RNA analysis community and help reduce the need for expensive IVT library preparation and sequencing for human samples. This strategy will serve as a framework for RNA modification analysis in other organisms.

## Methods

### Cell culture

HeLa, HepG2, A549, and NTERA-2 cells were cultured in Dulbecco’s modified Eagle’s medium (Gibco, 10566024) as a base; SH-SY5Y cells were cultured in a base of 1:1 EMEM:F12; Jurkat cells were cultured in RPMI (SH30027FS, FisherScientific). All media was supplemented with 10% Fetal Bovine Serum (FB12999102, FisherScientific) and 1% Penicillin-Streptomycin (Lonza,17602E). Cells were cultured at 37℃ with 5% CO_2_ in 10 cm tissue culture dishes until confluent.

### Total RNA extraction and Poly(A) selection

Total RNA extraction from cells and Poly(A) selection was performed using the protocol outlined previously[2]. Six 10 cm cell culture dishes with confluent cells were washed with ice-cold PBS and lysed with TRIzol (Invitrogen,15596026) at room temperature and transferred to an RNAse-free microcentrifuge tube. Chloroform was added to separate the total RNA in the aqueous supernatant from the organic phase containing DNA and cell debris below following centrifugation. The aqueous supernatant was then transferred to a fresh RNAse-free microcentrifuge tube, and an equal volume of 70% absolute ethanol was added. PureLink RNA Mini Kit (Invitrogen, 12183025) was used to purify the extracted total RNA in accordance with the Manufacturer’s protocol. Total RNA concentration was measured using the Qubit™ RNA High Sensitivity (HS) assay (Thermo Fisher, Q32852).

Poly(A) selection was performed using NEBNext Poly(A) mRNA Magnetic Isolation Module (NEB, E7490L) according to the Manufacturer’s protocol. The isolated Poly(A) selected RNA was eluted from the beads using Tris buffer. The poly(A) selected RNA concentration was measured using the same Qubit™ assay listed above.

### *In vitro* transcription and polyadenylation

The protocol for IVT, capping, and polyadenylation is described previously[2]. Briefly, the cDNA-PCR Sequencing Kit (SQK-PCS109) kit facilitated the reverse transcription (RT) and strand switching (SS). VN and Strand-Switching primers were added to 100 ng of poly(A) selected RNA from the abovementioned step. cDNA was produced by Maxima H Minus Reverse Transcriptase (Thermo Scientific, EP0751). Using a thermocycler, the reaction protocol is as follows: RT and SS for 90 minutes at 42°C (1 cycle), Heat inactivation for 5 mins at 85°C (1 cycle), and hold at 4°C (∞). PCR amplification was performed using LongAmp Taq 2X Master Mix (NEB, M0287S) and the Nanopore_T7_IVT_Forward and Reverse primers. The thermocycling conditions are as follows: initial denaturation for 30 seconds at 95°C (1 cycle), denaturation for 15 seconds at 95°C (11 cycles), annealing for 15 seconds at 62°C (11 cycles), extension for 15 minutes at 65°C (11 cycles), a final extension for 15 minutes at 65°C (1 cycle), and then an indefinite hold at 4°C. The PCR products were treated with Exonuclease 1 (NEB, M0293S) to digest single-stranded products. The resulting product was purified using Sera-Mag Select beads (Cytiva, 29343045) according to the Manufacturer’s protocol. IVT was performed on the purified PCR product using a HiScribe T7 High yield RNA Synthesis Kit (NEB, E2040S) and purified using a Monarch RNA Cleanup Kit (NEB, T2040S), both according to the Manufacturer’s protocols. Polyadenylation was performed using E. *coli* Poly(A) Polymerase (NEB, M0276S) using 12 μg of input RNA, and EDTA was added to halt the reaction. The product was purified again using a Monarch RNA Cleanup Kit with a final elution volume of 12 μL. Concentration was taken using the Qubit™ RNA HS assay. All primers are listed in Supplementary Table S1.

### Sequencing, Basecalling, and Alignment Procedure

Each cell line was sequenced individually using Nanopore Direct RNA sequencing (DRS) on R9 flow cells with sequencing chemistry SQK002. DRS runs were base called with Guppy v6.4.2 using the high accuracy model using the default basecalling quality score filter of Q ≥7[19]. Basecalled reads were aligned with minimap2(v2.24 [20]; RRID:SCR_018550) to the GRCh38.p10 reference genome and Gencode.v45 transcript sequences:

Gencode.v45: minimap2 -ax map-ont

GRCh38.p10: minimap2 -ax map-ont -uf -k 14

SAMs were filtered to include only primary alignments for downstream analysis: samtools view -h -F 4 -F 256 -F 2048

### NanoPlot[21]

Nanoplot was used to generate sequencing and gencode alignment stats: NanoPlot --raw

### NanoCount[22]

NanoCount (no options) was used to calculate transcript abundance. Basecalled reads were aligned to Gencode.v45 transcript sequences similarly as above with the additional option (-N 1) and not filtered further. HeLa biological DRS raw data was sourced from NIH NCBI-SRA BioProject: PRJNA777450

### Genomic DNA Extraction and Sanger Sequencing

We performed Sanger sequencing on HeLa genomic DNA (gDNA) to analyze putative mismatches. gDNA extraction was performed using a Monarch Genomic DNA Purification Kit (NEB, T3010S) following the Manufacturer’s protocol for cultured cells with an input of 5e6 HeLa cells. PCR primers to amplify ∼200 nt regions surrounding the mismatches were designed using Primer-BLAST with default settings and checking primer specificity against the *Homo Sapiens* genome. These primers are listed in Supplementary Table S1. Using the Manufacturer’s protocol, the PCR reaction was set up with Q5 polymerase (NEB, M0491L). Thermocycling conditions were as follows: initial denaturation at 98 ℃ for 30 seconds, 25 cycles of 98℃ for 10 seconds, then 63 ℃ for 20 seconds and 72 ℃ for 15 seconds, final extension at 72 ℃ for 2 minutes, and holding at 10 ℃. PCR products were purified using a Monarch PCR & DNA Cleanup Kit (NEB, T1030S) following the Manufacturer’s protocol. The concentration of eluted DNA was determined using a Nanodrop. The purified PCR products were imaged on a 2% agarose TBE gel to confirm specific amplification. Samples were sent to Quintara Biosciences for SimpliSeq^TM^ Sanger sequencing.

### Mismatch Analysis Methods

To identify the scope of mismatches in the sequenced IVT data (Supplementary Fig. S1), we followed an align, pileup, and compare strategy. We used the alignments to the GRCh38.p10 reference genome.

A positional pileup for each IVT replicate was created with pysamstats.

> pysamstats (--t variation)

The initial variant calling process documented every location with a minimum of 10x coverage in the pooled cell line data set with at least one called nucleotide differing from the reference. Subsequent filtering removed variants based on the number of occurrences of a given variant in relation to the number of canonical bases and deletions present at a location. We sequentially applied a filter of 30%, 40%, 50%, 60%, 70%, 80%, and 95% to the data.

Each set of variants was divided into three distinct bins: variants previously documented in Ensembl, variants that occurred in low-confidence 9mers, and novel variants for the IVT replicate set.

The final data set consisted of 3 compartments for each variant presence threshold: a set of potentially novel IVT variants, variants with known documentation in Ensembl, and variants that occurred in low-confidence 9mers (Supplementary Table S2).

### Round Robin Gene Coverage Saturation

To calculate the degree to which 5 of our cell lines (representative population) could approximate the observed gene set of the 6th, we subsampled 1,000,000 reads from the 6th cell line (target cell line) to create a representation of which that cell line covered genes. For each of the 5 other cell lines, we iteratively subsampled between 0 and 1,000,000 reads in increments of 100,000, for a combined total of 0 to 5,000,000 reads, and created a set of observed genes from those reads. At each iteration, we divided the cardinality of the union of our representative population with the target population by the cardinality of our target population to yield a proportion of the covered value. We repeated our sampling of the representative population 100 times and averaged the results for each sample interval. We repeated this process with each cell line acting as the target cell line. The iterative process was performed 500 times, yielding 6 saturation curves.

### Nanopolish Eventalign

Eventalign data for each dataset was computed with the following options nanopolish eventalign --print-read-names --scale-events --samples

### Data Validation and quality control

#### Generation and characterization of panIVT

We extracted RNA from 6 immortalized human cell lines: A549, HeLa, HepG2, Jurkat, NTERA, and SH-SY5Y cells and selected polyadenylated transcripts. These transcripts were reverse transcribed to cDNA, then *in vitro* transcribed back into RNA using canonical rNTPs according to Tavakoli et al. (see Methods). Each library was prepared for sequencing using an ONT Direct RNA Sequencing kit and sequenced on a MinION or PromethION flow cell. The throughput for these data ranged from 1.3 million to 4.9 million primary aligned reads per cell line (Table 1). Basecalling was performed using Guppy 6.4.2 and alignment using minimap2 to Gencode v.45 human reference transcripts (Fig. 1A)(Table 1).

**Figure 1.**
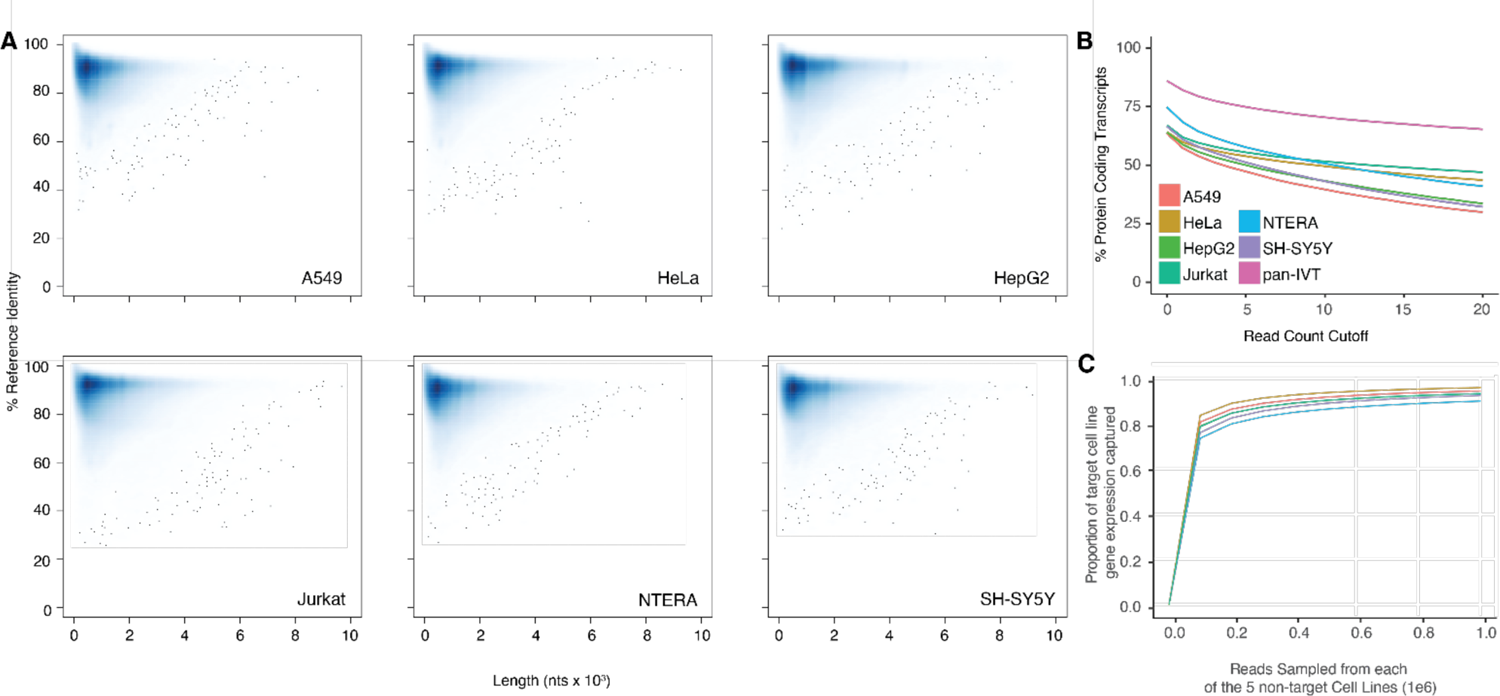
Alignment performance and coverage comparison. (**A**) Average percent identity of aligned DRS reads to Gencode.v38 Transcript sequences against read length (nts); data generated with Nanoplot[21]. (**B**) Percent of observed protein-coding transcripts (out of total protein-coding transcripts) against transcript minimum read count cut-off for each cell line. pan-IVT is the additive combination of all cell line coverage (**C**) Target cell line gene representation as a function of reads sampled from five sample population cell lines. Each cell line was used as the target cell line; the line on the graph corresponds to the representation of the target cell line listed in the legend.

**Table 1.**
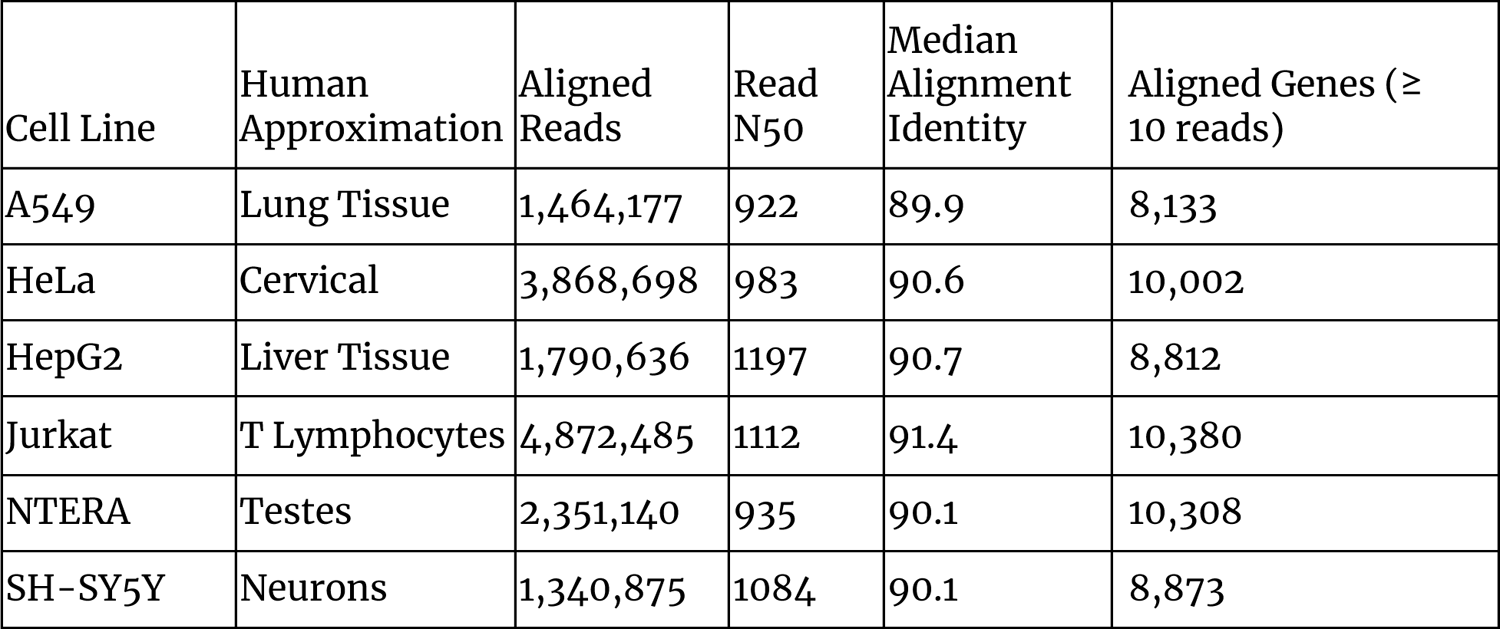
Cell Type Alignment Statistics.

We pooled the aligned IVT reads from each cell line (“panIVT”). We observed 17,038 unique genes aligned with at least one DRS read from the panIVT comprising 85.69% of all human protein-coding genes in Gencode v45. As the read count cutoff increased, the human protein-coding gene coverage decreased (Fig. 1B). As the panIVT comprises all six cell lines, it maximizes coverage and provides a more comprehensive representation of human protein-coding genes. We tested the ability of the five cell lines to capture the gene level diversity of the sixth cell line for increasing subsample sizes (Fig. 1C). NTERA has the lowest proportion of observed genes represented in the sample population at ∼90%. Combining different cell lines captured most of the observed gene level diversity for a cell line of interest.

### Cataloging sequence variation in IVT data using the GRCh38 reference

For RNA modification analysis, it is essential to distinguish systematic and spurious mismatches in IVT data. Systematic mismatches could be caused by genomic variation, IVT errors (i.e., due to reverse transcriptase or polymerase), sequencing-related errors, basecalling errors, or alignment errors. The sites that show these types of systematic errors in the IVT dataset should be omitted from RNA modification analysis. For each cell line, we identified positions with a mismatch percentage of 30% or higher from the GRCh38 reference genome. We further separated mismatches into one of three categories; known variants according to Ensembl[23], mismatches occurring in low confidence 9-mers (i.e., 9-mer regions where the basecaller is systematically less confident), and a combination of remaining mismatches (Supplementary Fig. S1; Table S2). The low confidence 9-mer set was curated by taking the average phred score base quality measure of all 9-mers in an independent biological DRS dataset[13] and selecting the lowest quartile. From our HeLa dataset with a minimum read count of 10 and mismatch occurrence threshold of 30%, we identified 62,708 mismatches. Of these 24,879 known variants, 8,930 originated in low-confidence 9-mers, leaving the remaining 28,899 mismatches (Supplementary Table S2). Without further orthogonal investigation, we recommend excluding all positions with a mismatch percentage at the preselected threshold, regardless of mismatch category. A tabulated version of the mismatch analysis findings for each cell line in our IVT dataset, as well as a pooled version, is publicly available on GitHub in conjunction with BED (Browser Extensible Data) files for each variance threshold for integrated genome viewing (IGV). The overlap of mismatches between each of the cell lines for the 30% cutoff threshold can be found in Supplementary Table S3.

### Re-use potential

#### Application of IVT dataset for downstream RNA modification analysis

We intend these data to be a negative control for biological DRS modification analysis. Transcript coverage and abundance correlation between a paired IVT mRNAs and its corresponding biological mRNA can assess the quality of the IVT as a negative control. For example, we compared the transcripts per million (TPM)[22] of IVT RNA from HeLa cells to biological mRNA from HeLa cells for aligned transcripts[2]. The TPMs were positively correlated (r^2^ = 0.83) indicating the IVT is representative of the biological data (Supplementary Fig. S2). Figure 2A details an example decision process to determine whether a candidate DRS site should be included in the downstream analysis using this IVT dataset. The mismatched BED files can serve as a first filtration point for sites where modification analysis is not recommended. These sites could include RNA editing sites, kmers that cause uncertainty in basecalling, or genetic variation that can’t be confirmed in the target sample. A stringent analysis would eliminate sites by using the 30% IVT mismatch filter. However, individual experiments may benefit from raising the occurrence threshold. We examined HeLa biological DRS data and BED files for 30%, 60%, and 95% occurrence thresholds in IGV (Supplementary Fig. S3A). Here, we can visualize positional nucleotide anomalies in HeLa biological DRS data. We can then refine the modification candidate sites list by excluding positions where an IVT mismatch occurs at a given occurrence threshold (Supplementary Fig. S3B). If a particular site is being examined where a reference mismatch occurs (either IVT or biological), we recommend using orthogonal methods to confirm IVT as an appropriate negative control at this position. This should include but isn’t limited to sequencing gDNA from the same sample as the direct RNA to confirm that the IVT identity and the DNA identity are the same. We recommend that biological conclusions drawn from analyses using these IVT data should include sufficient additional orthogonal confirmation.

**Figure 2.**
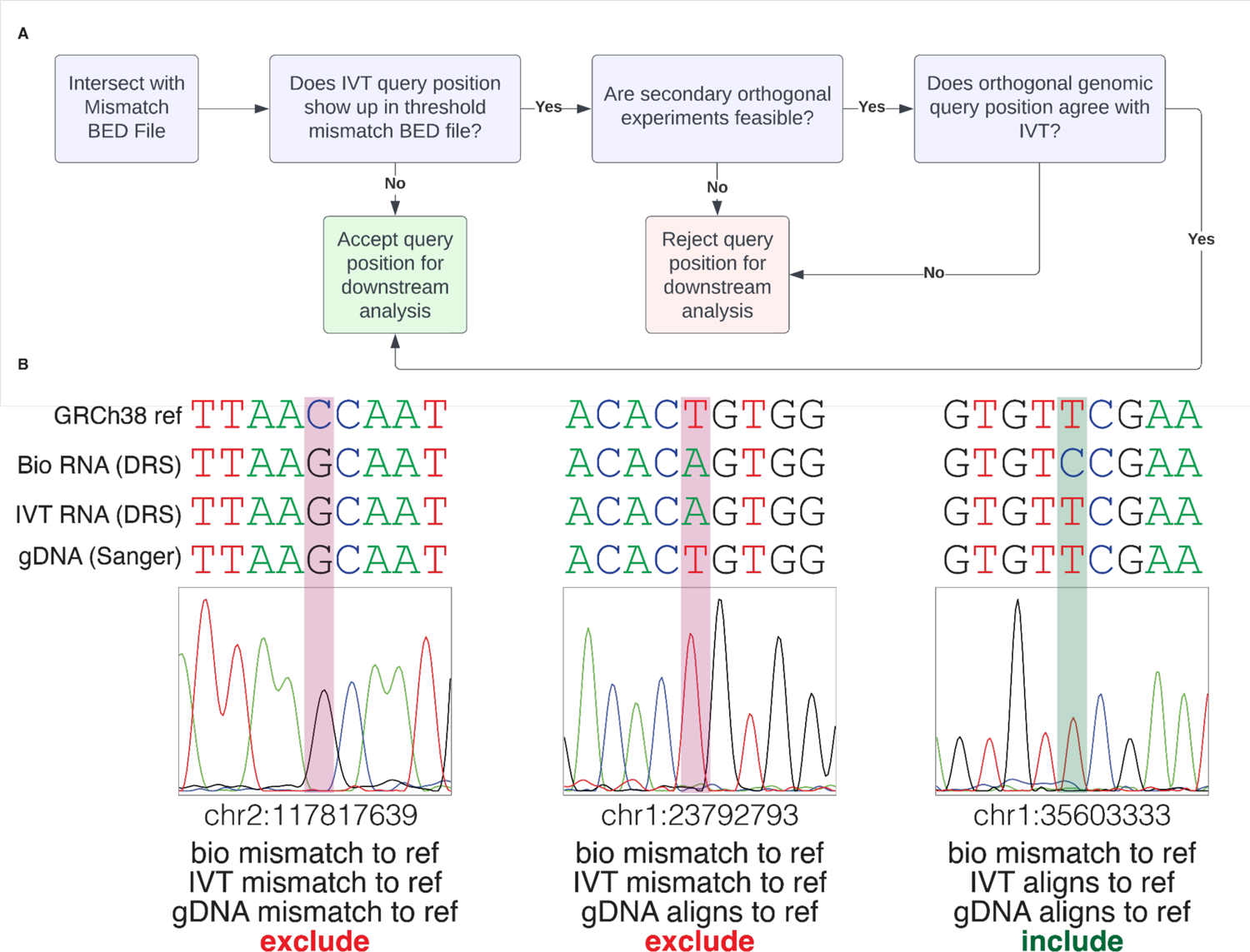
Recommended Analysis Inclusion Criteria (A) Decision tree to determine if a position should be considered for downstream analysis. (B) Sanger sequencing for orthogonal support determines suitability for downstream modification analysis. Comparing HeLa biological RNA (DRS), IVT RNA (DRS), gDNA (Sanger sequencing), and genome reference (GRCh38). Red bars indicate exclusion from downstream analysis, and green bars indicate inclusion.

We selected three example positions, determined their inclusion status (Fig. 2A), and performed Sanger sequencing[24] as an orthogonal method to confirm our decision (Fig. 2B). Two positions (chr2:117817639; chr1:23792793) show a mismatch between the observed IVT sequence and the reference sequence (at 80% occurrence threshold); we recommend the third position (chr1:35603333) as a candidate site for downstream modification analysis.

At the first position of interest (chr2:117817639), both the biological RNA and IVT RNA are mismatched to the reference (GRCh38.p10). Sanger sequencing at chr2:117817639 reveals the gDNA matches the biological and IVT RNA, indicating a single nucleotide variant (SNV). In this instance, we resolve to exclude this site as the biological RNA and IVT RNA agree, strongly suggesting the absence of an RNA modification. Again, the biological and IVT RNA mismatch to the reference nucleotide at the second position of interest (chr1:23792793). Unlike the preceding example, Sanger sequencing reveals the gDNA matches the reference, indicating the mismatch arose from a confounding variable. Therefore, this position is excluded from downstream analysis. The third position (chr1:35603333) represents a case where the IVT RNA matches the reference, but the biological RNA mismatches both the IVT RNA and the reference. Based on this information, we default to include this position for downstream analysis. Sanger sequencing confirms our decision, where the gDNA agrees with the IVT RNA and reference, indicating a candidate site for further RNA modification analysis.

Once the candidate positions are identified, various bioinformatic tools exist that leverage IVT as a negative control. Some tools, such as Mod-*p* ID[2], compare the rate of mismatches observed in biological and IVT data; for this application, the publicly available bam files can be used. Other tools, such as nanocompore[11] and xPore[12], require nanopolish eventalign data[19] computed from the raw sequencing files. Since eventalign is a computationally intensive program, we have precomputed the transcriptomic (gencode.v45) eventalign data and the genomic (GRCh38.p10) eventalign data and made them publicly available (https://github.com/RouhanifardLab/PanHumanIVT/tree/main). Figure 3A shows ionic current distributions for an example of five overlapping 5-mers. Cell lines with low coverage for the target 5mer had noisier ionic current distributions.

**Figure 3.**
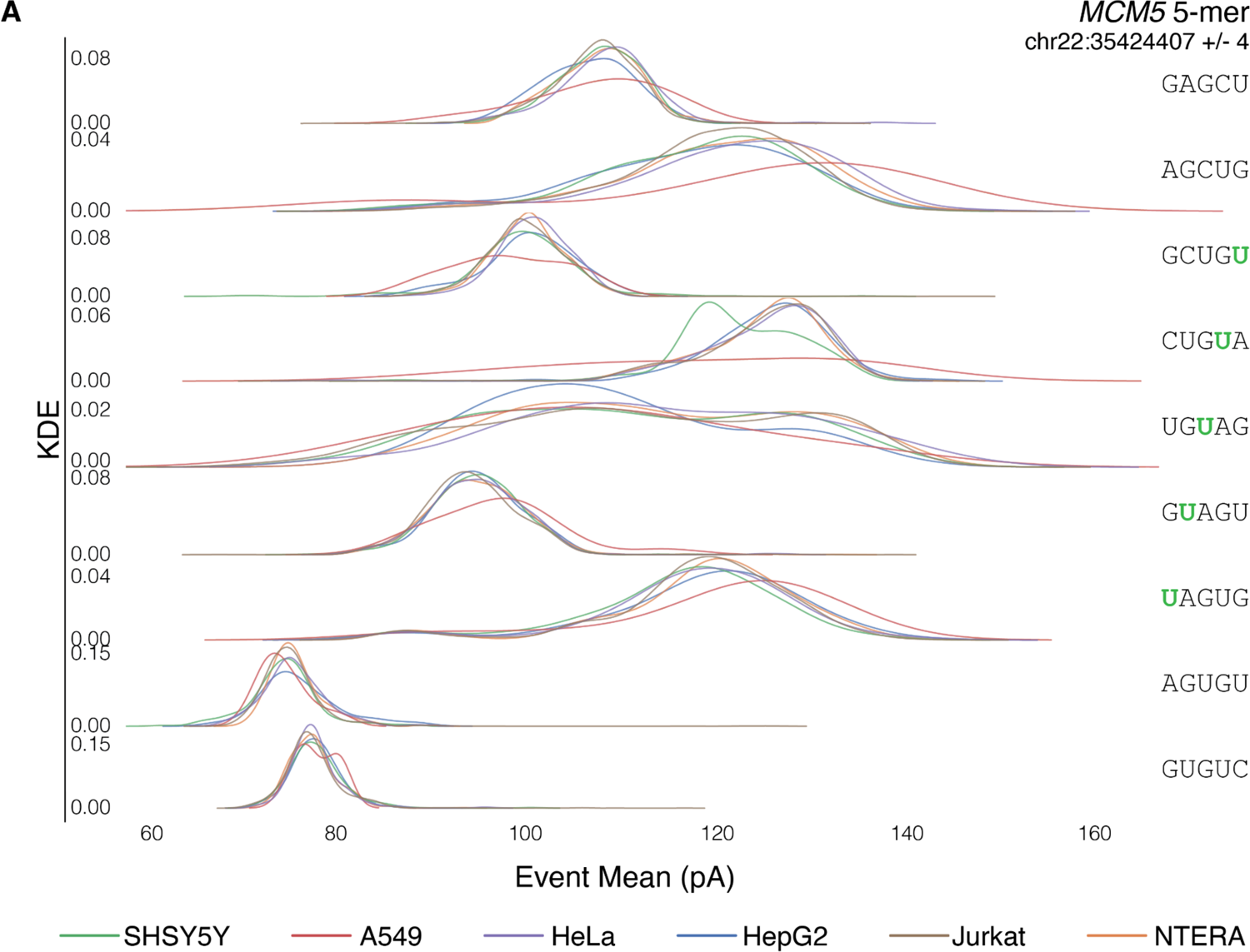
(A). Ionic Current distributions for *MCM5* (chr22:35424407 +/- 4) across all 6 cell lines.

## CONCLUSION

Appropriate negative controls are critical for accurately detecting and characterizing RNA modifications using nanopore DRS. IVT-derived negative controls provide an unmodified rendition of the transcriptome, allowing for comparative RNA modification analysis. A single cell line’s IVT transcripts may not fully capture the diversity of the human transcriptome, but expansion to multiple cell lines provides a more comprehensive representation.

We created IVT DRS data for six immortalized cell lines. We cataloged sites where the IVT dataset did not match the human reference genome. With this information, filtering out sites with potentially confounding underlying sequences and drawing more robust conclusions during comparative analysis is possible. The underlying sequence variation can come from several factors, including genomic variations introduced during the *in vitro* transcription process. Either source of error makes the position a poor candidate for comparative modification analysis. Because of this, we recommend eliminating IVT mismatch sites, regardless of origin, from comparative analysis.

Once the candidate sites have been selected, bioinformatic tools can be applied to perform comparative RNA modification analysis. We have precomputed and made publicly available nanopolish eventalign data to reduce the computational burden for potential users of this data set. While these tools can have high barriers of entry, we hope that this data set can help lower the computational burden and make RNA modification analysis more approachable for the community.

## Availability of source code and requirements

### Project name: PanHumanIVT

Project home page: https://github.com/RouhanifardLab/PanHumanIVT

DOI: https://doi.org/10.5281/zenodo.7976171

Operating systems: Linux Programming language: Python, R Other requirements: samtools 1.16.1 (using htslib 1.16), python 3.7, R 4.1.1, NanoPlot 1.40.2, Jupyterlab 3.4.4, NanoCount 1.0.0.post6 License: MIT License

## DATA AVAILABILITY

The data sets supporting the results of this article are available in the RouhanifardLab/PanHumanIVT GitHub repository, DOI: https://doi.org/10.5281/zenodo.7976171 FASTQ files and Fast5 raw data generated in this work have been made publicly available in NIH NCBI-SRA under the BioProject accession PRJNA947135 Sequences were aligned to genome version hg38.p10 and gencode version 45 transcript sequences available at: https://www.gencodegenes.org/

## ABBREVIATIONS

DRS: direct RNA sequencing

IVT: *in vitro* transcription

Ψ: pseudouridine

m^6^A: *N*^6^-methyladenosine

I: inosine

poly-A: polyadenylated

gDNA: genomic DNA

panIVT: pooled aligned IVT reads

BED: Browser Extensible Data

IGV: integrated genome viewing

SNV: single nucleotide variant

## AUTHOR CONTRIBUTIONS

C.A.M. and S.H.R. conceived of the research. C.A.M. and S.A. designed the experiments. C.A.M., S.T., D.B., and I.N.K. performed experiments and sequencing runs. C.A.M., S.A., D.B., and I.N.K. analyzed the data with guidance from S.H.R., and M.J.. C.A.M. and S.A. wrote the paper with guidance from S.H.R. and M.J.

## ADDITIONAL FILES

**Supplementary Fig. S1**. IVT to reference mismatch identification workflow.

**Supplementary Fig. S2**. mRNA coverage (TPM) correlation between HeLa IVT mRNA and HeLa biological mRNA.

**Supplementary Fig. S3**. IGV visualization of HeLa DRS and .bed files for 30, 60, and 95% SNV occurrence thresholds (OT). (A) view of chr2 displaying HeLa biological DRS read count depth (log scale) with corresponding IVT mismatch sites at each occurrence threshold. colors indicate mismatch presence for respective nucleotide (red = T, green = A, blue = C, orange = G) b, 100 nucleotide visualization of WDFY1 (chr2:223,877,264-223,877,362) where the reference nucleotide from hg38 genome is displayed as seq. Alignment mismatches for HeLa biological DRS are visualized proportionally as nucleotide count for respective colors. Grey indicates no significant mismatch at that position. Known variants at each occurrence threshold are denoted using the color of the variant nucleotide at that position.

**Supplementary Table. S1.** Primers used (*See Methods*)

**Supplementary Table. S2.** Cell Line IVT positional binned mismatch stats

**Supplementary Table. S3.** Overlapping mismatches between all cell lines for the 30% occurrence threshold.

