## Supplementary figures and images for "Multicellular, IVT-derived, unmodified human transcriptome for nanopore-direct RNA analysis"

### Supp Fig 1

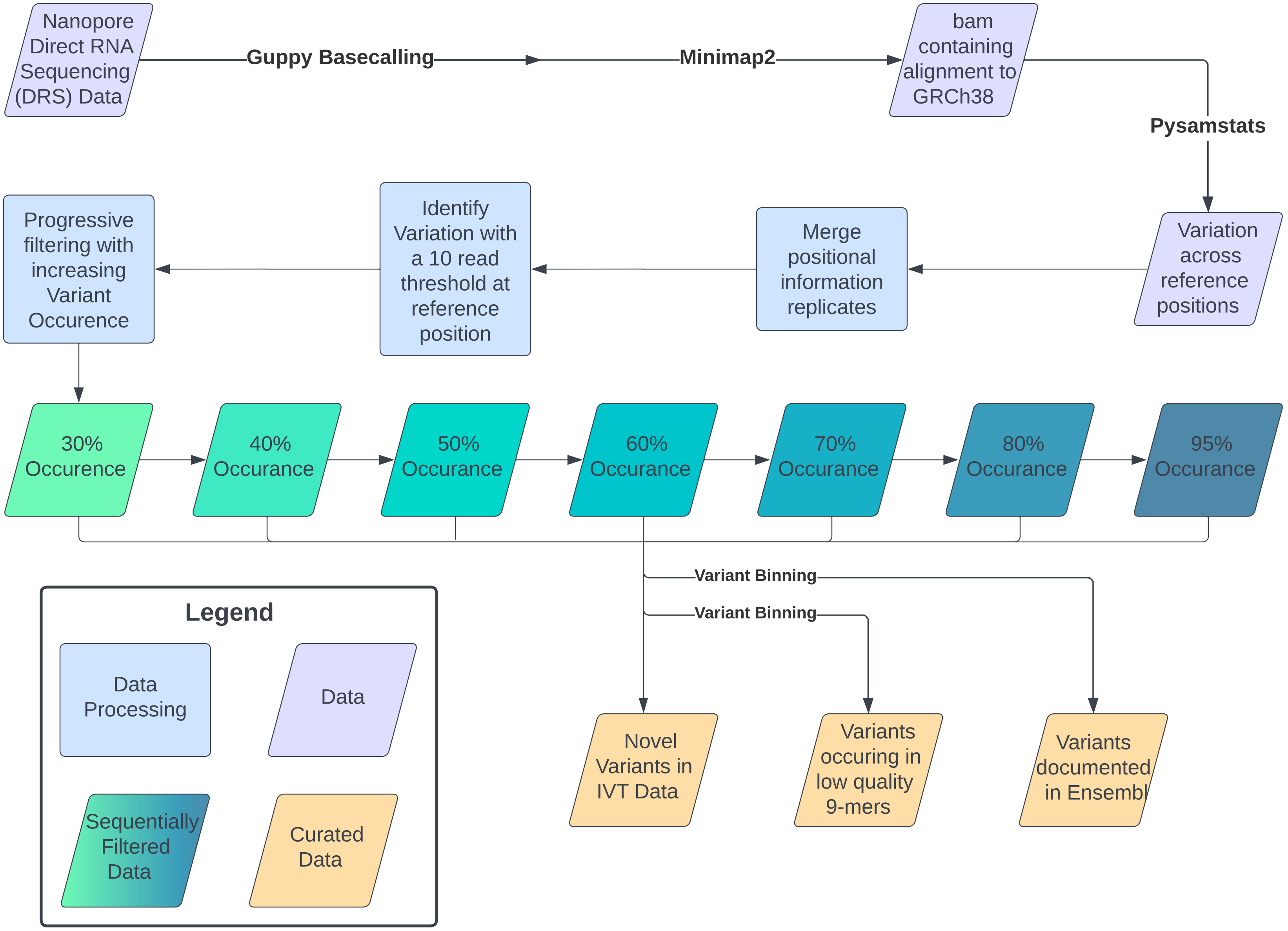

### Supp Fig 2

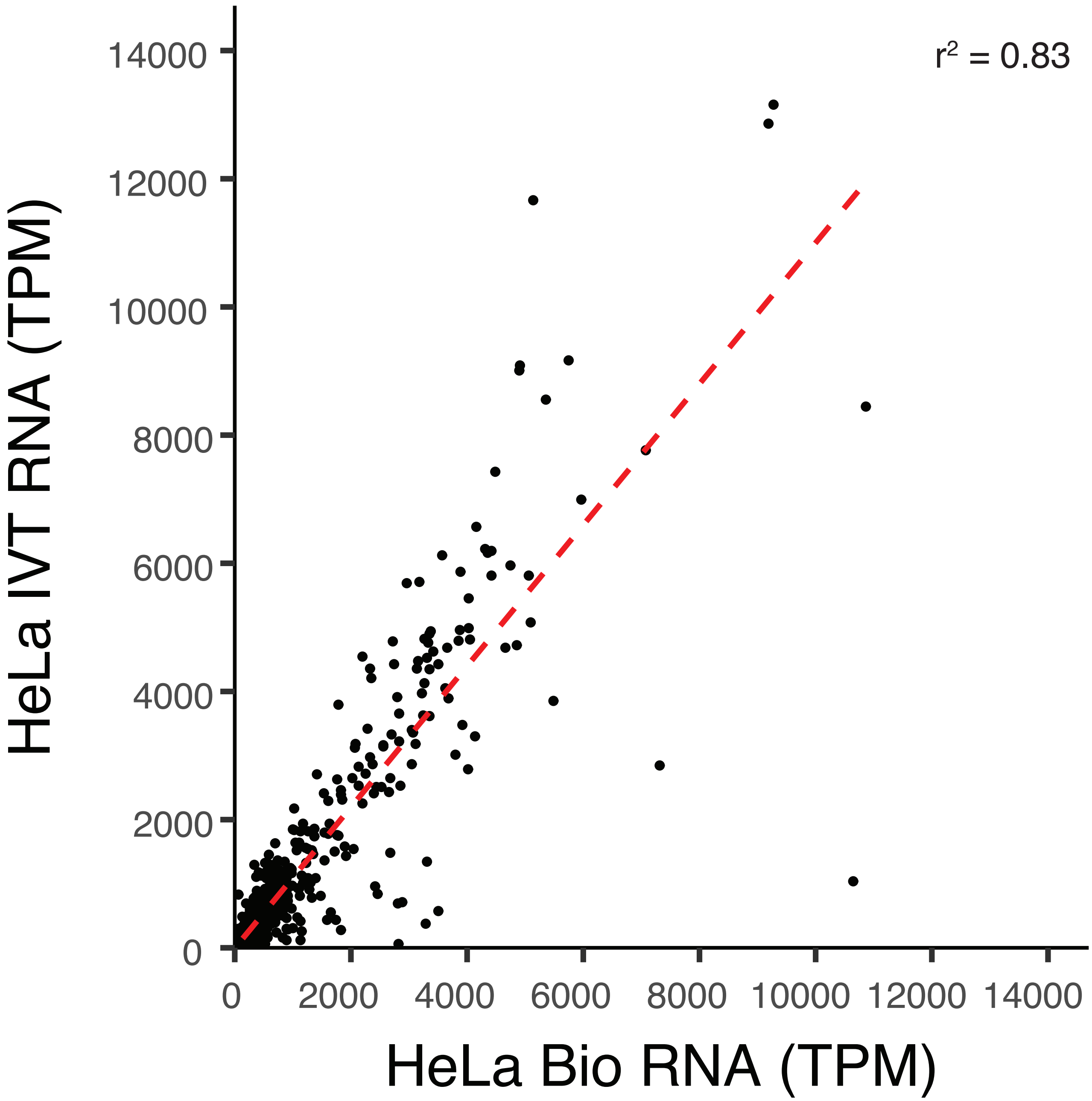

### Supp Fig 3

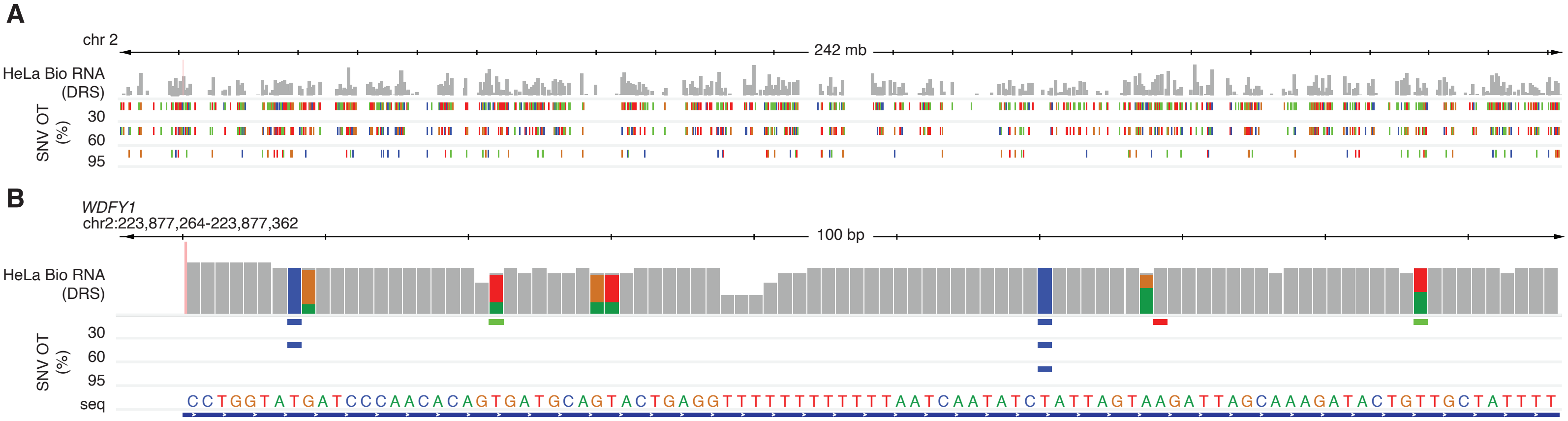
