## Supplementary material for "Multicellular, IVT-derived, unmodified human transcriptome for nanopore-direct RNA analysis": Supp Table 1

| Protocol Use | Oligo name | Sequence ( 5' -> 3') |
| --- | --- | --- |
| IVT Fwd Primer PCR | Nanopore_IVT_T7_Forward_Primer | TAATACGACTCACTATAGCGAGGCGGTTTTCTGTTGGTGCTGATATTGCT |
| IVT Rev Primer PCR | Nanopore_IVT_T7_Reverse_Primer | ACTTGCCTGTCGCTCTATCTTC |
| PCR F primer for sanger seq at Chr1:23792793 | Sanger_ Chr1:23792793_F | CCGTGTGGTGTATGTGTGGT |
| PCR R primer for sanger seq at Chr1:23792793 | Sanger_ Chr1:23792793_R | CAGGTAGCAGCCAAACAGGT |
| PCR F primer for sanger seq at Chr2:117817639 | Sanger_Chr2:117817639_F | GGAGGCATGTCTCAAGAAGCA |
| PCR R primer for sanger seq at Chr2:117817639 | Sanger_Chr2:117817639_R | AAACTAAATGGCTGAAGTTCAAAGA |
| PCR F primer for sanger seq at Chr19:2917188 | Sanger_Chr19:2917188_F | ACTGTGGACGAAAAGCACCT |
| PCR R primer for sanger seq at Chr19:2917188 | Sanger_Chr19:2917188_R | TCCGACACTGCTCGCATTT |
| PCR F primer for sanger seq at Chr3:19950940 | Sanger_Chr3:19950940_F | GGACATGGCTAGTCGAGGC |
| PCR R primer for sanger seq at Chr3:19950940 | Sanger_Chr3:19950940_R | AGAAAATCTCACCCCCAATGGT |
| PCR F primer for sanger seq at Chr4:109816233 | Sanger_Chr4:109816233_F | ATGTCTTTTCGAGGCGGAGG |
| PCR R primer for sanger seq at Chr4:109816233 | Sanger_Chr4:109816233_R | GGTCCTTGGTCTTGGCCTTT |
| PCR F primer for sanger seq at Chr1:35603333 | PSMB2_PseudoU_F_Primer | TGTTTGGGTACCCTCTACCAC |
| PCR F primer for sanger seq at Chr1:35603333 | PSMB2_PseudoU_R_Primer | AGGACATGATGTTAGGAGCCC |
