## Supplementary material for "Multicellular, IVT-derived, unmodified human transcriptome for nanopore-direct RNA analysis": Supp Table 2

| Cell Line | Occurrence Threshold | Bin | Quantity |
|---|---|---|---|
| HepG2 | 30% | Known Variants | 17962 |
| HepG2 | 30% | Low Confidence 9mers | 7300 |
| HepG2 | 30% | Novel Mismatches | 23430 |
| HepG2 | 40% | Known Variants | 9808 |
| HepG2 | 40% | Low Confidence 9mers | 1805 |
| HepG2 | 40% | Novel Mismatches | 5773 |
| HepG2 | 50% | Known Variants | 6424 |
| HepG2 | 50% | Low Confidence 9mers | 640 |
| HepG2 | 50% | Novel Mismatches | 2039 |
| HepG2 | 60% | Known Variants | 4865 |
| HepG2 | 60% | Low Confidence 9mers | 308 |
| HepG2 | 60% | Novel Mismatches | 1010 |
| HepG2 | 70% | Known Variants | 3655 |
| HepG2 | 70% | Low Confidence 9mers | 167 |
| HepG2 | 70% | Novel Mismatches | 518 |
| HepG2 | 80% | Known Variants | 2482 |
| HepG2 | 80% | Low Confidence 9mers | 81 |
| HepG2 | 80% | Novel Mismatches | 270 |
| HepG2 | 95% | Known Variants | 535 |
| HepG2 | 95% | Low Confidence 9mers | 9 |
| HepG2 | 95% | Novel Mismatches | 40 |
| SH-SY5Y | 30% | Known Variants | 18715 |
| SH-SY5Y | 30% | Low Confidence 9mers | 8524 |
| SH-SY5Y | 30% | Novel Mismatches | 28762 |
| SH-SY5Y | 40% | Known Variants | 9596 |
| SH-SY5Y | 40% | Low Confidence 9mers | 2085 |
| SH-SY5Y | 40% | Novel Mismatches | 6903 |
| SH-SY5Y | 50% | Known Variants | 6067 |
| SH-SY5Y | 50% | Low Confidence 9mers | 698 |
| SH-SY5Y | 50% | Novel Mismatches | 2170 |
| SH-SY5Y | 60% | Known Variants | 4449 |
| SH-SY5Y | 60% | Low Confidence 9mers | 312 |
| SH-SY5Y | 60% | Novel Mismatches | 947 |
| SH-SY5Y | 70% | Known Variants | 3276 |
| SH-SY5Y | 70% | Low Confidence 9mers | 145 |
| SH-SY5Y | 70% | Novel Mismatches | 482 |
| SH-SY5Y | 80% | Known Variants | 2219 |
| SH-SY5Y | 80% | Low Confidence 9mers | 64 |
| SH-SY5Y | 80% | Novel Mismatches | 233 |
| SH-SY5Y | 95% | Known Variants | 431 |
| SH-SY5Y | 95% | Low Confidence 9mers | 8 |
| SH-SY5Y | 95% | Novel Mismatches | 36 |
| A549 | 30% | Known Variants | 13460 |
| A549 | 30% | Low Confidence 9mers | 5807 |
| A549 | 30% | Novel Mismatches | 19596 |
| A549 | 40% | Known Variants | 7231 |
| A549 | 40% | Low Confidence 9mers | 1436 |

| | | | |
|---|---|---|---|
| A549 | 40% | Novel Mismatches | 4900 |
| A549 | 50% | Known Variants | 5098 |
| A549 | 50% | Low Confidence 9mers | 474 |
| A549 | 50% | Novel Mismatches | 1753 |
| A549 | 60% | Known Variants | 3898 |
| A549 | 60% | Low Confidence 9mers | 234 |
| A549 | 60% | Novel Mismatches | 905 |
| A549 | 70% | Known Variants | 2884 |
| A549 | 70% | Low Confidence 9mers | 121 |
| A549 | 70% | Novel Mismatches | 523 |
| A549 | 80% | Known Variants | 1940 |
| A549 | 80% | Low Confidence 9mers | 60 |
| A549 | 80% | Novel Mismatches | 258 |
| A549 | 95% | Known Variants | 412 |
| A549 | 95% | Low Confidence 9mers | 7 |
| A549 | 95% | Novel Mismatches | 36 |
| NTERA | 30% | Known Variants | 20228 |
| NTERA | 30% | Low Confidence 9mers | 8983 |
| NTERA | 30% | Novel Mismatches | 29461 |
| NTERA | 40% | Known Variants | 11124 |
| NTERA | 40% | Low Confidence 9mers | 2259 |
| NTERA | 40% | Novel Mismatches | 7629 |
| NTERA | 50% | Known Variants | 7951 |
| NTERA | 50% | Low Confidence 9mers | 821 |
| NTERA | 50% | Novel Mismatches | 2875 |
| NTERA | 60% | Known Variants | 6158 |
| NTERA | 60% | Low Confidence 9mers | 399 |
| NTERA | 60% | Novel Mismatches | 1493 |
| NTERA | 70% | Known Variants | 4493 |
| NTERA | 70% | Low Confidence 9mers | 209 |
| NTERA | 70% | Novel Mismatches | 834 |
| NTERA | 80% | Known Variants | 3033 |
| NTERA | 80% | Low Confidence 9mers | 107 |
| NTERA | 80% | Novel Mismatches | 422 |
| NTERA | 95% | Known Variants | 581 |
| NTERA | 95% | Low Confidence 9mers | 14 |
| NTERA | 95% | Novel Mismatches | 46 |
| Jurkat | 30% | Known Variants | 31598 |
| Jurkat | 30% | Low Confidence 9mers | 12119 |
| Jurkat | 30% | Novel Mismatches | 37593 |
| Jurkat | 40% | Known Variants | 19864 |
| Jurkat | 40% | Low Confidence 9mers | 3567 |
| Jurkat | 40% | Novel Mismatches | 12143 |
| Jurkat | 50% | Known Variants | 13186 |
| Jurkat | 50% | Low Confidence 9mers | 1542 |

| | | | |
|---|---|---|---|
| Jurkat | 50% | Novel Mismatches | 5825 |
| Jurkat | 60% | Known Variants | 9854 |
| Jurkat | 60% | Low Confidence 9mers | 938 |
| Jurkat | 60% | Novel Mismatches | 3509 |
| Jurkat | 70% | Known Variants | 7600 |
| Jurkat | 70% | Low Confidence 9mers | 588 |
| Jurkat | 70% | Novel Mismatches | 2258 |
| Jurkat | 80% | Known Variants | 5355 |
| Jurkat | 80% | Low Confidence 9mers | 337 |
| Jurkat | 80% | Novel Mismatches | 1351 |
| Jurkat | 95% | Known Variants | 1167 |
| Jurkat | 95% | Low Confidence 9mers | 80 |
| Jurkat | 95% | Novel Mismatches | 280 |
