## Supplementary material for "Multicellular, IVT-derived, unmodified human transcriptome for nanopore-direct RNA analysis": Supp Table 3

| | Unique | In 1 Other | In 2 Others | In 3 Others | In 4 Others | In All |
|---|---|---|---|---|---|---|
| A549 mismatches | 18,733 | 5,121 | 3,986 | 3,119 | 3,091 | 3,813 |
| HeLa mismatches | 32,776 | 11,524 | 6,772 | 4,314 | 3,509 | 3,813 |
| HepG2 mismatches | 24,577 | 8,214 | 5,110 | 3,724 | 3,254 | 3,813 |
| Jurkat mismatches | 54,698 | 10,616 | 5,908 | 3,511 | 2,764 | 3,813 |
| Ntera mismatches | 30,492 | 9,996 | 6,536 | 4,264 | 3,571 | 3,813 |
| SH-SY5Y mismatches | 30,163 | 9,445 | 5,405 | 3,844 | 3,331 | 3,813 |
